# Environmental and geographic drivers of global bat phylogenetic diversity

**DOI:** 10.1101/2025.02.18.636314

**Authors:** Abigail E. Green, Camilo A. Calderón-Acevedo, J. Angel Soto-Centeno, Tara A. Pelletier

**Affiliations:** Department of Mathematics & Statistics, Radford University, Radford, VA 24142; Department of Biology, Radford University, Radford, VA 24142; Smithsonian Tropical Research Institute, Balboa, Ancón, Panamá; Department of Mammalogy, Division of Vertebrate Zoology, American Museum of Natural History, New York, NY 10024

**Author notes:** **Biosketch** Our team is interested in exploring biodiversity patterns in a wide-range of taxa using methods from population genetics, phylogeography, and systematics. AEG: www.linkedin.com/in/abigail-abbi-green TAP: https://sites.google.com/site/taraapelletier/ CCA: https://mammalbiogeo.wordpress.com/ ASC: https://www.mormoops.com/.

**Keywords:** Chiroptera, phylogenetic diversity, COI, species delimitation, ecoregions, random forest

## Abstract

**Aim:** Understanding the patterns and factors that shape biodiversity is vital to conserving species. We combined open-source genetic, environmental, and geographic information to analyze bat phylogenetic diversity (PD) patterns in continuous ecoregions across the globe. This information is important for developing bat conservation strategies, and our methodology can work for any taxa with sufficient georeferenced genetic data available.

**Location:** Global.

**Methods:** After curating a global dataset containing 14,037 COI DNA sequences from 343 described species of bats, we calculated PD for continuous ecoregions at different spatial scales. To avoid the difficulties of using current species names, we used genetic OTUs identified by a single-locus species delimitation method to reconstruct and date a phylogeny. We then calculated PD, estimated a lineage through time plot, and used random forest predictive modeling to identify environmental and geographic predictors of PD.

**Results:** In addition to current temperature, temperature during the last glacial maximum and temperature changes between the last glacial maximum and last interglacial were most closely associated with PD. However, at different spatial scales, the top variables differed slightly. When using smaller ecoregions, latitude and population density were also identified as important, though not significant. Southeast Asia and South America had the highest levels of PD, along with parts of Africa and the Himalayas. We demonstrate that, regardless of spatial scale and uneven sampling across the globe, single-locus genetic data can reflect species diversity gradients and identify predictors of PD.

**Main conclusions:** We show that publicly-available, single-locus data can be used to analyze large-scale evolutionary patterns and inform conservation efforts. Additionally, choice of biodiversity measure and spatial scale matter when assessing species patterns. When looking at bats, temperature variables with a historical component are most important for predicting PD broadly, but latitude and population density could also be important on smaller scales.

## Introduction

Considering multiple components of biodiversity is necessary for understanding the relative roles of evolutionary processes in shaping communities at different spatial scales (Stevens & Gavilanez, 2015), and different measures of biodiversity may vary across different spatial scales. For example, species richness and within-species genetic diversity are not always evenly or similarly distributed across geographic space and can be influenced by different evolutionary processes (Amador, et al., 2024; Brown et al., 2020; Lino, et al., 2021). Within-species genetic diversity and phylogeographic breaks follow a similar pattern to the latitudinal species diversity gradient (da Silva Fonseca, et al., 2024; Smith et al., 2014) while patterns of isolation-by-distance or -environment are better explained by range size (Pelletier & Carstens, 2018; Amador et al. 2025). Documenting these differing biodiversity components is important for structuring conservation efforts that are effective and efficient (Harper et al., 2021; Hosegood et al., 2020). Phylogenetic diversity accounts for branch lengths on a phylogenetic tree (Faith, 1992) or genetic distance (i.e., divergence) among species and is therefore a useful measure of biodiversity to consider for conservation purposes. Assemblages of species that contain more distantly related species have been shown to consist of and produce higher levels of biomass (Cadotte, 2013). More distantly related groups also capture more diversity in species traits, which increases future potential for adaptation (Forest et al., 2007) and may better preserve natural processes. Phylogenetic diversity is also listed as an important consideration by the International Union for Conservation of Nature (www.pdtf.org) and globally can contribute to the Convention on Biological Diversity’s global conservation efforts (www.cbd.int).

Most data available in open-source repositories can lead to new ways of documenting and exploring biodiversity, especially by integrating environmental, geographic, and genetic data. For instance, efforts to create global maps of biodiversity can be useful for tracking changes over time (Perreira 2016), identifying areas of high diversity in need of conservation (Pelletier, et al., 2018), and understanding the evolutionary processes promoting biodiversity (Guralnick & Hill, 2009; Theissinger et al., 2023). Recent studies using open-source data to document biodiversity provide information to facilitate conservation action and may help unravel the processes that shape highly speciose regions (e.g., phylogeographic breaks in birds: Smith, et al., 2017; response to Pleistocene glaciations in bats: Carstens, et al., 2018; hidden diversity in mammals: Parsons, et al., 2022). Conservation initiatives that include more complex levels of biodiversity, incorporate evolutionary history, and capture functional diversity can help maintain the stability of a wide range of taxonomic groups and minimize species loss. For example, when space for protected areas is limited, targeting phylogenetic diversity can more effectively conserve a larger proportion of the mammal tree of life (Rosauer, et al., 2017; Rosauer et al., 2018).

Increased anthropogenic land use and habitat fragmentation can negatively affect the composition of biodiverse communities (Newbold et al, 2015; López⍰Baucells et al., 2022). Habitat degradation reduces ecosystem stability and productivity which endangers important ecosystem services, like food production, at local and regional scales (Cardinale et al., 2012; Moreira-Hernández, et al., 2021). Additionally, the increased rate of the rise in global temperatures caused by human emissions of greenhouse gases and pollution is making many habitats inhospitable by humans, wildlife, and vegetation alike (Grimm et al., 2013; Weiskopf et al., 2020). A better understanding of the key environmental and geographic factors that both foster and discourage long-term biodiversity trends can help improve conservation decision making and land use management. Strong conservation initiatives are now more pertinent than ever given the rapid increase in biodiversity and habitat loss that has led to the endangerment and extinction of many species in recent millennia (Ceballos, et al., 2020; Newbold et al, 2015). Furthermore, climate change is inevitable and understanding how biodiversity responds to the climate in its environment can have useful conservation implications.

Many studies have examined the effects of environmental factors on phylogenetic diversity (PD) across multiple taxonomic and geographic scales (Grimshaw & Higgins, 2017; Carvalho, et al., 2021; Paz et al., 2021; Montaño-Centellas, et al., 2023; Hu et al., 2021). The importance of different environmental factors may change with the measure of phylogenetic diversity used, taxonomic group, or spatial scale (Grimshaw & Higgins, 2017; Paz et al., 2021; Montaño-Centellas et al., 2023). For example, Paz et al. (2021) found that, in plants, mean nearest taxon distance (MNTD) and mean pairwise distance (MPD) were more closely associated with temperature while Faith’s index was more closely associated with precipitation. Nonetheless, there is limited information about the effects of environmental factors on biodiversity globally because most studies are conducted at regional geographic scales (Hu et al., 2021). PD in the bats of Mexico (Grimshaw & Higgins, 2017) and the Atlantic Forest of Brazil (Stevens and Gavilanez 2015) was best explained by temperature, fluctuations in temperature, and latitude. Elevation predicted phylogenetic turnover in phyllostomid bats in the Amazon and Atlantic Forest of Brazil (de Carvalho et al., 2019; Carvalho et al., 2023). For all mammals in western Mexico, however, precipitation was a more important factor for taxonomic diversity (Mason-Romo et al., 2017). On a global scale, raptor PD was lower in areas with historically high temperatures and higher human disturbance (Montaño-Centellas et al., 2023).

Bats (order Chiroptera) comprise 1500 species, representing about 22% of crown mammal diversity (Mammal Diversity Database, 2024; Simmons & Cirranello, 2025). Two suborders and 21 families of bats are currently recognized (Teeling et al., 2005). The phylogenetic history of bats reflects a deep diversification event where two suborders diverged ∼63mya (Foley et al. 2023; Hao et al. 2024). The suborder Yinpterochiroptera is composed of five families of megabats and smaller insectivore and carnivore species. The suborder Yangochiroptera contains the remaining families and exhibits the largest ecological variation. This ecological variation is evident when comparing the most diverse families: Vespertilionidae, a largely insectivorous lineage with one carnivorous representative; Phyllostomidae in the Neotropics with less than half the bat species (230), but with the largest Eltonian and Grinnelian niche amplitudes of any mammal clade (Fleming et al., 2020); and Pteropodidae, the flying foxes, which are primarily nectarivorous and frigivorous. Given their unique ability to fly, many bats may be able to disperse across large areas providing important ecosystem services like insect pest control, pollination of over 500 species of angiosperms by the three aforementioned mentioned families (Muchhala & Tschapka 2020; Moréira-Hernández et al., 2021), and seed dispersal (Kunz, et al., 2011; Moréira-Hernández et al., 2021; Ramírez-Fráncel et al., 2022). Over one-third of bat species assessed by the IUCN are considered threatened or data deficient, and 80% have unknown or decreasing population trends (Frick, et al., 2020). Human derived activities at local, regional, and global scales are the main source of bat species decline and population fragmentation (Frick et al., 2020; Soto-Centeno & Calderón-Acevedo, 2023); however, we have limited knowledge on how phylogenetic diversity relates to habitat and environmental change across space and time in most taxonomic groups, especially at a global scale.

We explored how environmental and geographic factors affect bat phylogenetic diversity on a global scale. First, we estimated phylogenetic diversity using georeferenced DNA sequences and species delimitation methods across ecoregions. Then, we used a random forest predictive algorithm to determine which geographic and environmental variables had the largest impact on phylogenetic diversity measures in continuous ecoregions across the globe. We also included historical climatic data starting in the Pliocene because paleoclimate leaves impressions on species ranges and diversity (Svenning et al. 2015), so it will be important to consider future climate change in conservation planning on a global scale (i.e., ecoregions; Dobrowski et al. 2021). We expect largely heterogeneous ecoregions with more stable, warmer temperatures to contain the highest levels of PD given that landscape barriers to dispersal and time should promote speciation, particularly in tropical areas (Smith et al. 2014). Furthermore, given regional studies conducted in bats, climate variables pertaining to temperature and historical climate could also have an impact on PD on larger scales. Our study serves as a first step in defining areas that contain high levels of phylogenetic diversity and the conditions under which biodiversity could be maintained and promoted moving forward through estimates of PD across ecoregions with heterogeneous habitats that should capture broad-scale biodiversity and evolutionary potential.

## Methods

### DNA sequence data and OTUs

We obtained data from public databases, such as Genbank and the Global Biodiversity Information Facility (GBIF), through the *phylogatR* platform (Pelletier et al. 2022; https://phylogatr.org/) using the search term ‘Chiroptera’ on February 22, 2022. The total dataset represented 17 out of 21 (80%) families, 429 out of 1482 (29%) species, and 18 loci with a total of 20,020 DNA sequences. *PhylogatR* provides DNA sequence alignments on a species-by-species basis that are almost analysis ready. In this study, we focused on the mitochondrial gene *Cytochrome oxidase I* (*COI*) as this locus was the best represented in the data. We first checked for potentially misidentified sequences. Each species sequence alignment was run through BLAST via the NCBI database. The top hit for each DNA accession was compared to the species name associated with the DNA sequence in *phylogatR*. Five hundred thirty sequences that were considered not plausible data (i.e., too short) and two that did not BLAST to bats (i.e. *Mus musculus* and *Apodemus draco*) were removed from the alignments. Cleaned alignments from this step were merged for all species and re-aligned using MAFFT v7.5 (Kotah & Standley, 2013) with the default settings and –adjustdirection. We visually inspected the concatenated *COI* sequence alignment and trimmed the ends of the sequences to remove large areas with missing data resulting in a 658 base-pair region of this barcoding gene. Our final dataset consisted of 14,037 *COI* DNA sequences found globally (Fig. 1; Table S1). Of these, 1215 (8%) had species names in GenBank that did not match records existing in the GBIF or the Barcode of Life Data System (BOLD). Because we used OTUs as units of analysis, we chose to retain these taxa without adjusting taxonomy (details below).

**Figure 1:**
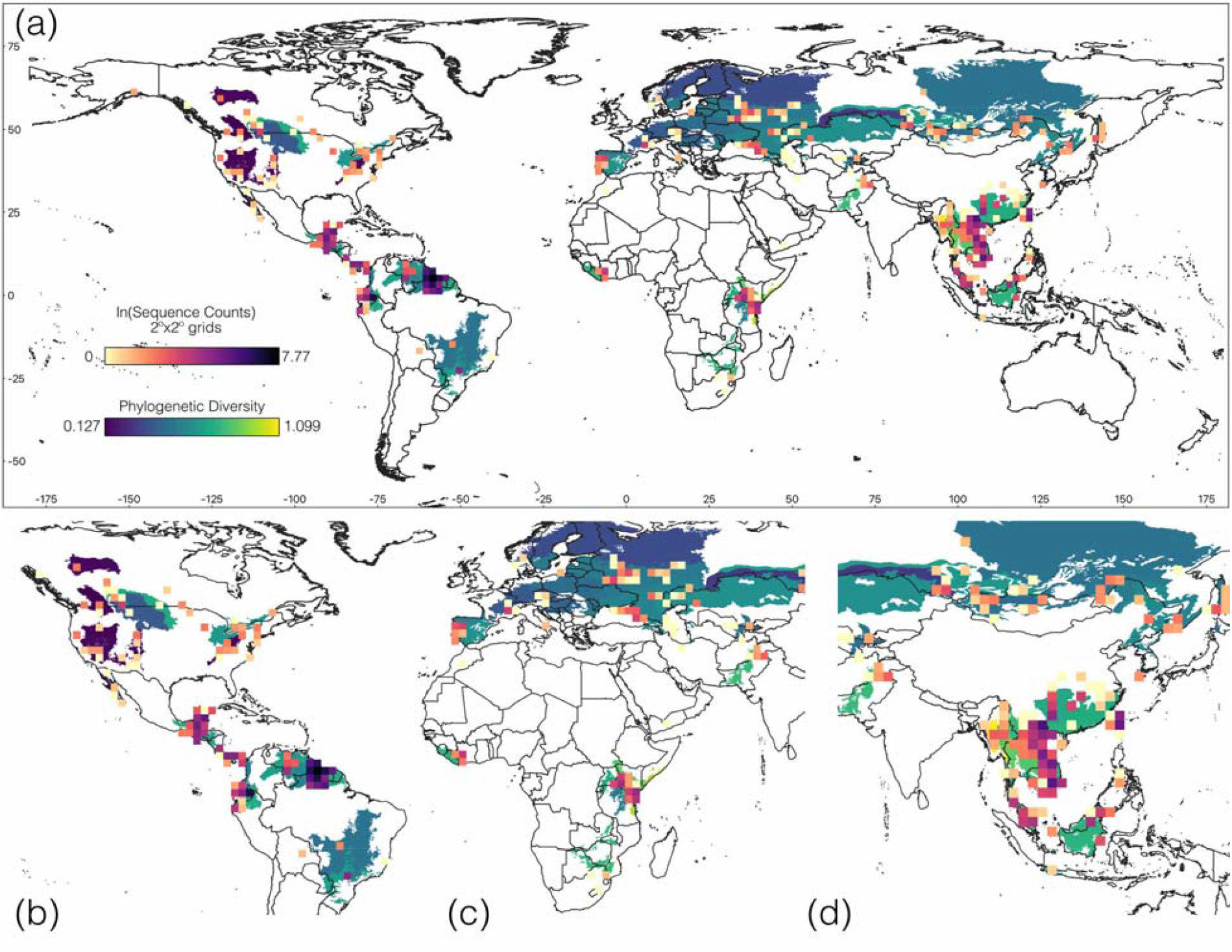
109-ecoregions. (a) Phylogenetic diversity of bats Worldwide and number of DNA sequences (given in natural logarithm of actual sequence numbers in a 2x2 degree grid, for better visualization). (b) The Neotropics (c) East Africa (d) Southeast Asia

Mammal taxonomy is rapidly changing (Burgin, et al., 2018; Parsons et al., 2022; Reeder, et al., 2007; Solari & Martínez-Arias, 2014; Simmons & Cirranello, 2025; Mammal Diversity Database, 2024) and species names are often incorrect in online databases (Mulcahy et al., 2022). Therefore, we used OTUs estimated from the DNA sequence data itself, rather than species names, as terminal taxa in our phylogeny. By using OTUs, we captured biodiversity above and below currently named species diversity, which can have useful conservation implications (Rosauer et al., 2018). This is crucial, particularly for groups with complicated taxonomy, such as bats (Calderón-Acevedo, et al., 2021; Calderón-Acevedo, et al., 2022; da Silva Fonseca, 2024; Mônico & Soto-Centeno, 2024; Rodríguez-Posada, et al., 2021; Yi et al., 2024) and small mammals in general (Parsons et al. 2022). To estimate genetic lineages and define OTUs, we used the distance-based species delimitation method Assemble Species by Automatic Partitioning (ASAP; Puillandre, et al., 2020), as tree-based methods (e.g., GMYC [Pons et al., 2006] or PTP [Zhang, et al., 2013]) would be prohibitive given the size of our dataset. All default settings in ASAP were used. This method establishes a set of ten different partitions ranked by an ASAP score. Each partition is a list of genetically distinct groups defining which individuals belong to each group. The ASAP score is a combination of the probability that each ASAP group is a single species along with the width of the barcode gap between the current and previous partition. Generally, the smaller the ASAP score, the better the partition is (Puillandre et al., 2020), so we used the partition with the lowest score (i.e., best ranked).

### Phylogeny reconstruction and estimates of divergence

One DNA sequence was randomly sampled from each ASAP genetic group (i.e., OTU) to estimate a phylogenetic tree representing each *COI* lineage present in our dataset. This phylogeny (Supporting files S1 & S2) was then used in subsequent estimates of divergence times, lineage through time plots, and phylogenetic diversity (see below). We first used ModelFinder (Kalyaanamoorthy et al., 2017) to select the best model of nucleotide evolution. Then, we implemented the resulting model (GTR+F+R8) in RAxML-NG (Kozlov, et al., 2019). This analysis ran for an initial search of 10 Maximum Likelihood (ML) trees followed by a 1000 bootstrap tree search (see Supporting Information). In this analysis, we used Yinpterochiroptera as an outgroup to all other bats following well-supported previously published bat relationships that combined mitochondrial and nuclear sequences (Shi & Raboski, 2015; Teeling et al., 2005). Given the limited information in a single-locus dataset, we wanted to ensure that we followed major accepted bat relationships (Teeling et al., 2005; Shi & Raboski, 2015; Baker, et al., 2016; Cirranello, et al., 2016; Rojas, et al., 2016). A configuration file (Supporting file S3) for the RaxML-NG analysis included constrained relationships at the level of superfamily and family, but allowing the inference of subfamily, genera, and species/OTU (all intermediate tree construction outputs can be found in Supporting files S1-S12).

We used the rooted constrained RaxML-NG phylogeny (Supporting files S2 and S9) to compute divergence time estimates with the penalized likelihood semi-parametric method available in TreePL (Smith & O’Meara, 2012). This method allows for the estimation of different rates on different branches on a previously estimated phylogeny with branch lengths. To estimate divergences, we used the node ages of Yingochiroptera and Yangochiroptera (71–58 my), Noctilionoidea (37–47 my), the node of Pteropodidae and Rhinolophoidea (53–63 my), the node between *Pteronotus* and *Mormoops* (36.2–30.8 my), the internal node of Stenodermatinae and Phyllostominae within Phyllostomidae (36.2–27.33), and the first node within the subfamily Desmodontinae (23.9–16.7 my) as calibration points previously available from the literature (see Teeling et al. 2005; Shi & Raboski, 2015; Rojas et al., 2016). The TreePL analysis then followed an initial priming run where optimization parameters were obtained, followed by a cross-validation step to select the best smoothing parameter for the penalized likelihood estimates (Supporting file S10, Smith & O’Meara, 2012). The resulting divergence time estimates (Supporting file S11) were used to construct a lineage through time (LTT) plot with the R packages ‘ape’ and ‘phytools’ (Paradis & Schliep, 2019; Revell, 2024). The LTT plot was produced using the family Phyllostomidae as a reference because this clade of bats shows a remarkable rate of species diversification in the Neotropics (Shi & Raboski, 2015; Rojas et al., 2016).

### Phylogenetic diversity and geographic units, ecoregions

The most common ways to measure phylogenetic diversity include Faith’s index, mean pairwise distance (MPD), and mean nearest taxon distance (MNTD) (Faith, 1992; Warwick & Clarke, 1995; Webb, et al., 2002; Webb, 2000). Faith’s index sums of all the branch lengths in a phylogenetic tree. Mean pairwise distance calculates average distance among all species in a phylogenetic tree. Mean nearest taxon distance calculates the average distance between a species and its closest evolutionary neighbor. Because MPD and MNTD include measurements of branch lengths based on nearest neighbors and not total branch lengths, these measures, MPD in particular, should be somewhat protected from missing species, as they focus on relationships between tips of the phylogeny (Cadotte et al. 2012; Li et al. 2012); however, we used the standardized effect size (SES) approach for MPD and MNTD to better control for missing taxa. OTU richness was calculated for each ecoregion (see below), and all three measures of PD were estimated using the R package ‘picante’ (Kembel et al., 2010).

To estimate levels of PD across the globe, we chose to analyze data by ecoregion rather than using grid cells that may be arbitrary from a biological perspective (Paz-Vinas et al., 2021). From a conservation perspective, an ecoregion approach can yield higher representation of novel habitats, natural communities, and species-level diversity than other units of analysis such as protected areas or political boundaries; therefore, more adequately protecting species, their natural processes, connectivity, and the species they interact with (Funk and Fa 2010; Giakoumi et al. 2013; Hanson et al. 2022; Zoderer et al. 2024). Our goal was not to focus on single species or endemics, but to use the presence or absence of a lineage in each ecoregion to identify the ecoregions that may foster the persistence of biodiversity in bats, as ecoregions are ecologically cohesive but span a suite of novel habitats and specialized niches (Zoderer et al. 2024).

We used the 846 RESOLVE Ecoregions 2017 dataset (Olson et al., 2001; Dinerstein et al., 2017). There were 161 ecoregions containing DNA sequences in our dataset. However, many of those had very few species, limiting our ability to make comparisons of OTU level diversity. For example, the 161 ecoregions ranged from 2 – 85 OTUs per ecoregion and had an average of 8.96 OTUs per region. There were 109 out of 161 ecoregions that had more than two OTUs represented and could be used to estimate PD. We also created a second set of larger continuous ecoregions by grouping RESOLVE Ecoregions that contained DNA sequences from our dataset, were adjacent to each other, and belonged to the same biome. This dataset contained 25 continuous ecoregions, from 185 RESOLVE Ecoregions, as some regions contained small fragmenting ecoregions that were included to create a continuous space (Table S2). There were 3 – 117 OTUs per ecoregion with an average of 22.44 OTUs. We used both datasets, a 109-eco set that used the smaller single ecoregions and a 25-eco set with combined ecoregions, to conduct subsequent analyses. Investigating the same sets of predictor variables at different spatial scales allowed us to determine whether spatial scale can influence trends in bat diversity.

We also conducted linear regressions to test whether OTU richness explained the different measures of PD to determine whether PD captures a different feature of biodiversity other than simply OTU richness, especially given that our bat dataset is not complete (i.e., open-source databases rarely contain all species). OTU richness explained almost all of the variation in Faith’s index (109-eco: R^2^ = 0.9767, p < 0.001; 25-eco: R^2^ = 0.9815, p < 0.001), so we moved forward with predictive modeling on PD measurements that were not significantly correlated with OTU richness: MNTD (109-eco: R^2^ = 0.04689, p = 0.02372; 25-eco: R^2^ = 0.03237, p = 0.1925) and MPD (109-eco: R^2^ = 0.1139, p < 0.001; 25-eco: R^2^ = 0.1218, p = 0.0488). PD estimates were used as the response variable in the predictive modeling framework described below.

### Predictor variables

We curated a set of predictor variables using 115 environmental data layers (Table S3) and geographic information associated with each ecoregion. We took the median and standard deviation of each data layer in each region by randomly sampling 1000 points per region. This was done to capture information about the ecoregion as a whole and the habitat heterogeneity that contributes to or inhibits connectivity, which might promote or hinder speciation and be important under future climate change scenarios (Dobrowski et al. 2021). Identifying the environmental factors that contribute to biodiversity may help predict areas that could harbor the most biodiversity. The data layers included 19 BIOCLIM layers for present time from the Chelsa database (Karger et al., 2017; Karger et al., 2018; Karger, et al., 2021) at 1km resolution, elevation (Tachikawa et al., 2011), population density (CIESIN, 2017), terrestrial habitat heterogeneity (Tuanmu & Jetz, 2015), gross domestic product (GIS processing World Bank DECRG, 2010), global land cover classification (European Space Agency, 2009), global river classification (Dallaire, et al., 2019), disaster risk (Peduzzi, 2019), anthropogenic biome (Ellis, et al., 2010), global lakes and wetlands database (Lehner & Döll, 2004), and various indicators of seasonal growth (Karger et al., 2021). We also included BIOCLIM layers for four different historical time periods: Mid-Holocene (ca. 8.326-4.2 ka); Last Glacial Maximum (LGM, ca. 21 ka), Last Interglacial (LIG, ca. 130 ka), and Pliocene (ca. 3.3 ma) (Fordham et al., 2017; Karger et al., 2021; Otto-Bliesner, et al., 2006; Dolan, et al., 2015). We used the R packages ‘raster’ (v3.6-14; Hijmans, 2023), ‘tools’ (R Core Team, 2022), and ‘sf’ (Pebesma, 2018) to extract ecoregion specific information for each of these layers. We also calculated the median, standard deviation, maximum, and minimum values for latitude and longitude of each ecoregion and used the st_area function to calculate the area of each ecoregion (Pebesma, 2018).

To capture change of climate variables over time, we calculated the difference in the median values between time periods: Pliocene—LIG, LIG—LGM, LGM—Holocene, and Holocene—present in addition to the median and standard deviation of each layer. This resulted in 310 variables that represent a variety of environmental and geographic traits for each ecoregion (Tables S4 & S5). Twelve of the 310 variables for the 25-eco dataset contained NA values and were removed from the dataset, as the random forest algorithm does not handle missing data. These twelve variables included the median and standard deviation for three growing season temperature variables, snow water equivalent, and two global lakes and wetlands classifications. There were six additional variables that contained NA values and were removed for the 109-eco dataset: the mean and standard deviation for the last day of the growing season, the first day of the growing season, and global drought events. The resulting 25-eco dataset contained 298 variables and the 109-eco dataset contained 292 variables for further exploration.

### Predictive modeling

To reduce the number of variables in our predictive models, we used different sets of predictor variables based on correlation coefficients and variable categories. We first calculated pairwise Pearson correlation coefficients for all environmental and geographic variables using the R package ‘Hmisc’ (Harrell Jr., 2023) and removed predictor variables with a correlation coefficient greater than or equal to 0.75 (Tables S6 & S7). We then split the variables into seven groups based on category of variable and ran a Random Forest (RF) for each group: 1) current climate, 2) geography (including latitude, longitude, flood risk, growing season, anthropogenic biomes, etc.), 3) Holocene climate, 4) LGM climate, 5) LIG climate, 6) Pliocene climate, and 7) climate change (the difference across time periods). We ran these category analyses using all variables and with the correlation cutoff of 0.75. In total, we ran 14 RF analyses with different sets of predictor variables for both MPD and MNTD as response variables with the R package ‘randomForest’ (Liaw & Wiener, 2002) using the default settings, 1000 trees, and retaining the importance values. While random forest handles a high variable to observation ratio relatively well (Genuer et al. 2008), we conducted 100 iterations for each RF to assess stability of the model.

Based on the predictive power of the RF models (Table 1), we focus on MPD as our phylogenetic diversity response variable. Percent increase in mean squared error (MSE) was calculated for each variable when removed from the model to determine which variables contribute the most to the prediction. We used this value to demonstrate the relative importance of each variable. To further reduce the variables in our data and assess which were most important, we took the top 2-4 predictors from the seven random forest models (each category listed above with 0.75 or higher correlated variables removed) to run a final RF. Variables were chosen for the final RF if they were ranked in the top five most important predictors for their model in 100% of the RF iterations . There were 19 top predictors for the 109-eco set and 23 top predictors for the 25-eco set across the seven different categories of variables (Tables S8 & 9). For the top predictors in the final RF models, we used the R package ‘nlme’ (Pinheiro & Bates, 2000; Pinheiro et al., 2025) to conduct generalized least squares (GLS) linear regressions and test the overall significance of the predictor variables while accounting for the spatial autocorrelation that commonly arises in geographical analyses like ours.

**Table 1.**
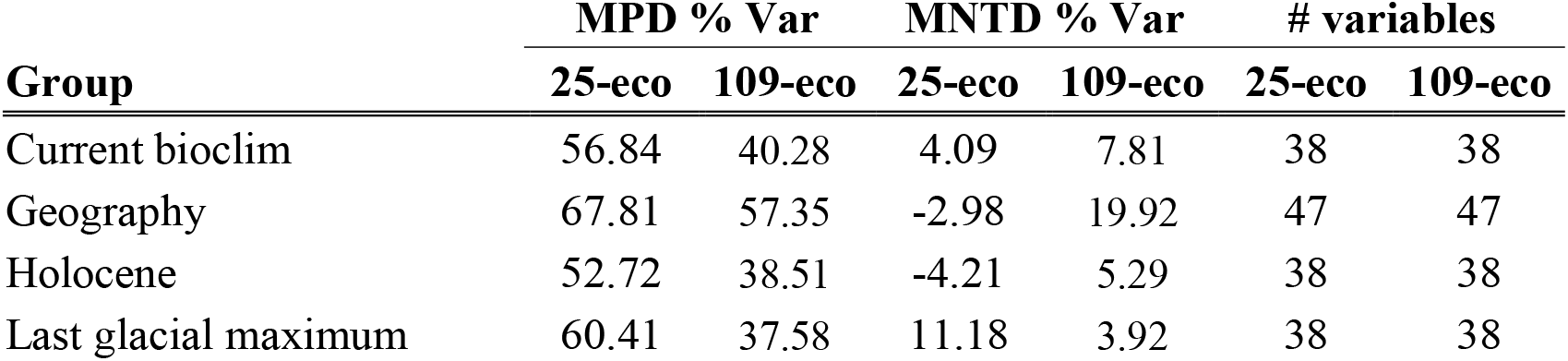
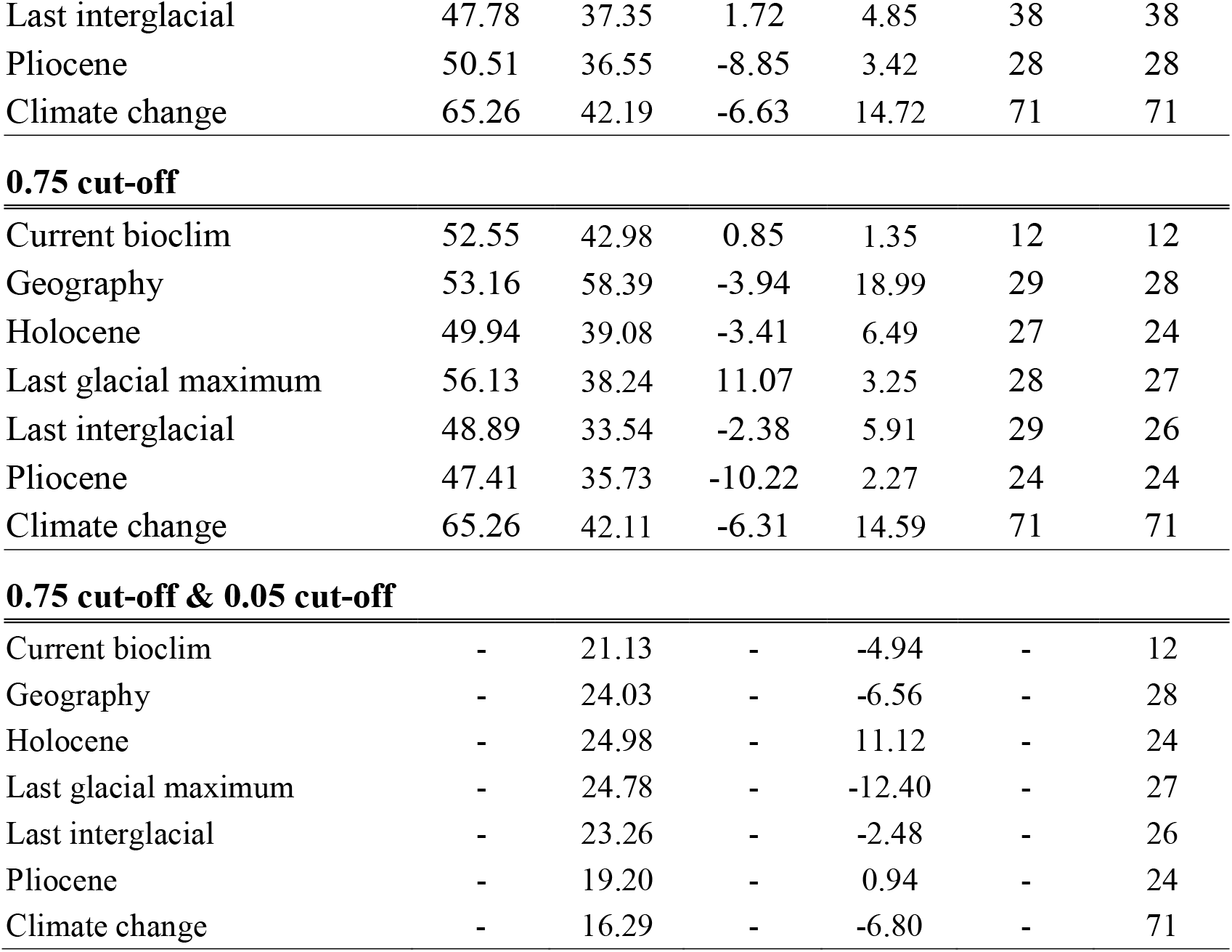
Percent variation explained for all random forest models and the number of predictor variables used in each model. 0.75 cut-off refers to the variable set where those correlated higher than 0.75 were removed. 0.05 cut-off refers to the set where ecoregions were removed if their PD values were not significant after randomization. BIOCLIM variables for present day (Current), the Holocene, the Last Glacial Maxima (LGM), the Last Interglacial (LIG), and the Pliocene (Plio) were grouped into their own categories. The final two categories were geographic variables (Geography), which also contained other data layers, and Climate change which included differences between each of time periods for all BIOCLIM variables.

| Group | MPD % Var |  | MNTD % Var |  | # variables |  |
| --- | --- | --- | --- | --- | --- | --- |
|  | 25-eco | 109-eco | 25-eco | 109-eco | 25-eco | 109-eco |
| Current bioclim | 56.84 | 40.28 | 4.09 | 7.81 | 38 | 38 |
| Geography | 67.81 | 57.35 | -2.98 | 19.92 | 47 | 47 |
| Holocene | 52.72 | 38.51 | -4.21 | 5.29 | 38 | 38 |
| Last glacial maximum | 60.41 | 37.58 | 11.18 | 3.92 | 38 | 38 |
| Last interglacial | 47.78 | 37.35 | 1.72 | 4.85 | 38 | 38 |
| Pliocene | 50.51 | 36.55 | -8.85 | 3.42 | 28 | 28 |
| Climate change | 65.26 | 42.19 | -6.63 | 14.72 | 71 | 71 |
| 0.75 cut-off |  |  |  |  |  |  |
| Current bioclim | 52.55 | 42.98 | 0.85 | 1.35 | 12 | 12 |
| Geography | 53.16 | 58.39 | -3.94 | 18.99 | 29 | 28 |
| Holocene | 49.94 | 39.08 | -3.41 | 6.49 | 27 | 24 |
| Last glacial maximum | 56.13 | 38.24 | 11.07 | 3.25 | 28 | 27 |
| Last interglacial | 48.89 | 33.54 | -2.38 | 5.91 | 29 | 26 |
| Pliocene | 47.41 | 35.73 | -10.22 | 2.27 | 24 | 24 |
| Climate change | 65.26 | 42.11 | -6.31 | 14.59 | 71 | 71 |
| 0.75 cut-off & 0.05 cut-off |  |  |  |  |  |  |
| Current bioclim | - | 21.13 | - | -4.94 | - | 12 |
| Geography | - | 24.03 | - | -6.56 | - | 28 |
| Holocene | - | 24.98 | - | 11.12 | - | 24 |
| Last glacial maximum | - | 24.78 | - | -12.40 | - | 27 |
| Last interglacial | - | 23.26 | - | -2.48 | - | 26 |
| Pliocene | - | 19.20 | - | 0.94 | - | 24 |
| Climate change | - | 16.29 | - | -6.80 | - | 71 |

Finally, we conducted another round of RF with the 109-eco set regions that had significant values for the SES MPD randomization procedure (Table S5). Given that the original dataset contained missing taxa, we wanted to ensure that any signals in the data were consistent and reliable. This dataset contained 41 ecoregions. We only used predictor variables sets with highly correlated variables removed because we were starting with a smaller number of observations. We also only conducted these analyses with MPD as the prior models with MNTD as the response variable did not demonstrate any predictive power. We followed the same procedure as above and retained only the top variables from each category for a final RF analysis (Table S10), followed by GLS to assess significance.

## Results

### Genetic data and OTUs

After data cleaning and trimming the sequence alignment, there were 658 base-pairs of *COI* from 343 species and 14,037 individual sequences (Fig. 1; Table S1). ASAP identified 372 genetic partitions or OTUs. With >14,000 sequences, the ASAP analysis took less than 20 minutes on iMac (3.6 GHz 8-Core Intel Core i9 16 GB 2667 MHz DDR4). The ASAP results largely matched species names but there were cases where nominal species were split, merged, and sometimes both (Fig. S1; Table S11). More specifically, 155 nominal bat species matched the ASAP groups exactly, where all individual sequences from each species only belonged to one ASAP group, 81 nominal species were split into multiple ASAP groups, and 161 nominal species were combined in 69 ASAP groups. A total of 54 nominal species that were split into multiple ASAP groups also had individuals merged with other nominal species. The average value for OTU richness across the 25-eco set was 21.48 with East Asia having the highest (116) and four different regions (Mountainous region of Asia (Himalayas), Northwest Asia, Northwest North America, and the Philippines) having the lowest (3) (Table S4). The average value for OTU richness across the 109-eco set was 12.28 with the Guianan lowland moist forests having the highest (85), followed by the Northern Annamites rain forests (53), and 23 regions with the lowest (3) (Table S5).

### Bat phylogeny reconstruction and lineages through time

The estimated phylogeny (Fig. 2 & S2; Supporting file S2) accurately inferred most of the currently accepted relationships of bats both at the levels of subfamily and genus. Estimates of lineages through time based on the reconstructed phylogeny demonstrated a pattern congruent with rapid species diversification in Phyllostomidae, the most speciose Neotropical bat family, compared to more gradual lineage accumulation for other more deeply or recently divergent families (e.g., Molossidae and Hipposideridae; Fig. 3 & S3). However, we noted that some deep nodes (i.e., the sister relationship of Emballonuroidea with and Noctilionoidea) in the phylogeny were recovered with short branches (Fig. 2). Furthermore, the divergence time estimates produced by TreePL show deeper (i.e., older) divergences than expected based on previously published studies (e.g., Teeling et al., 2005; Rojas et al., 2016). These observations were likely due to our use of a single mitochondrial marker with a fast mutation rate compared to other studies that use a combination of nuclear and mitochondrial data. Despite that, our choice of using a single mitochondrial marker allowed us to include a wide sampling breadth across 70% of all bat families worldwide, which facilitated the analysis of drivers of diversity globally (read below).

**Figure 2:**
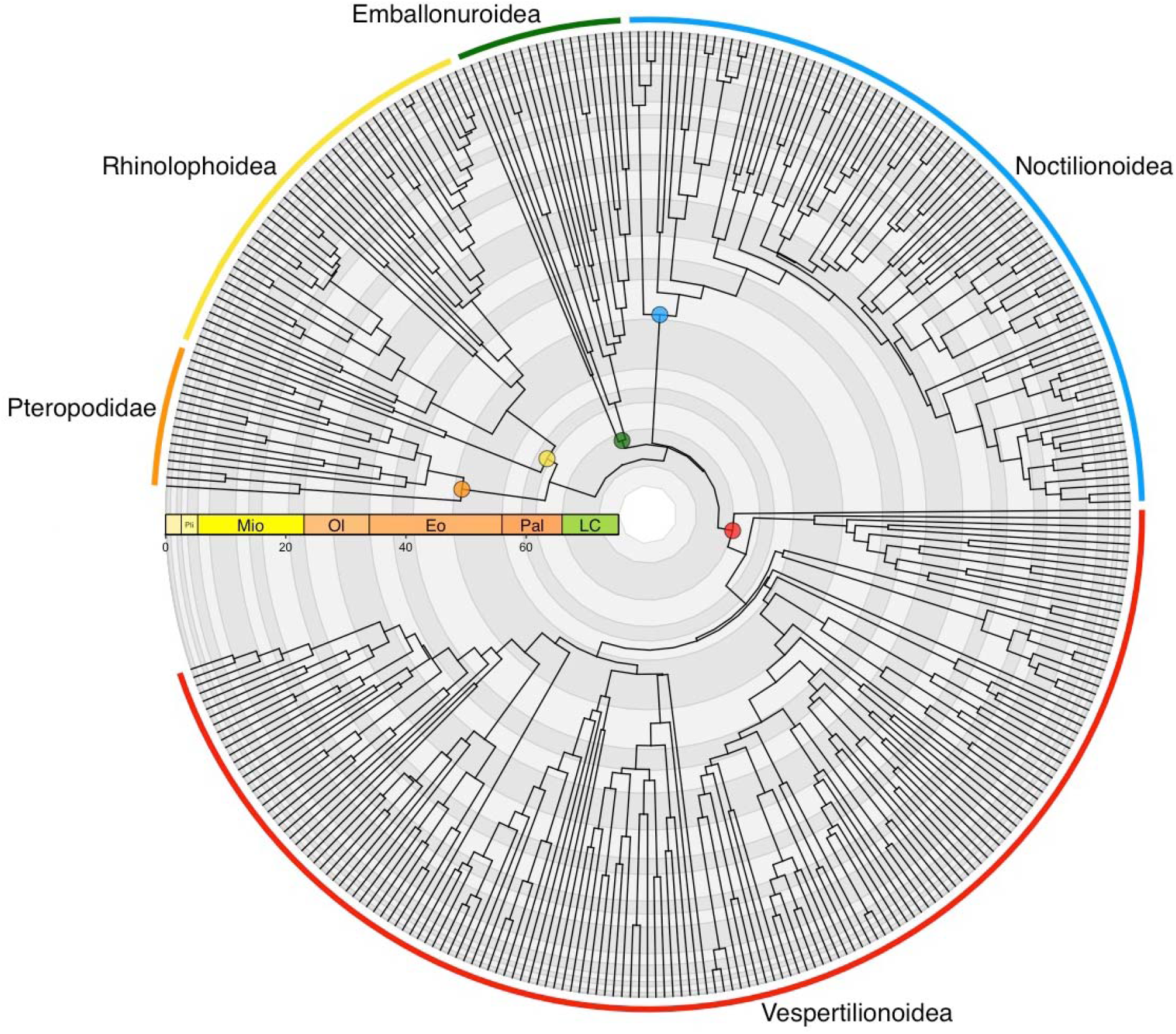
Time calibrated phylogeny of bats worldwide representing 17 families and all extant superfamilies. Estimates of divergence were performed in TreePL using the RaxML-NG COI phylogeny. Color coded nodes represent the superfamily and family taxonomic designations at tips of the phylogeny. Time scale represented in millions of years and following the geological time designations Pal= Paleogene, Eo = Eocene, Ol = Oligocene, Mio = Miocene, and Pli = Pliocene (the Pleistocene and Holocene not labeled).

**Figure 3:**
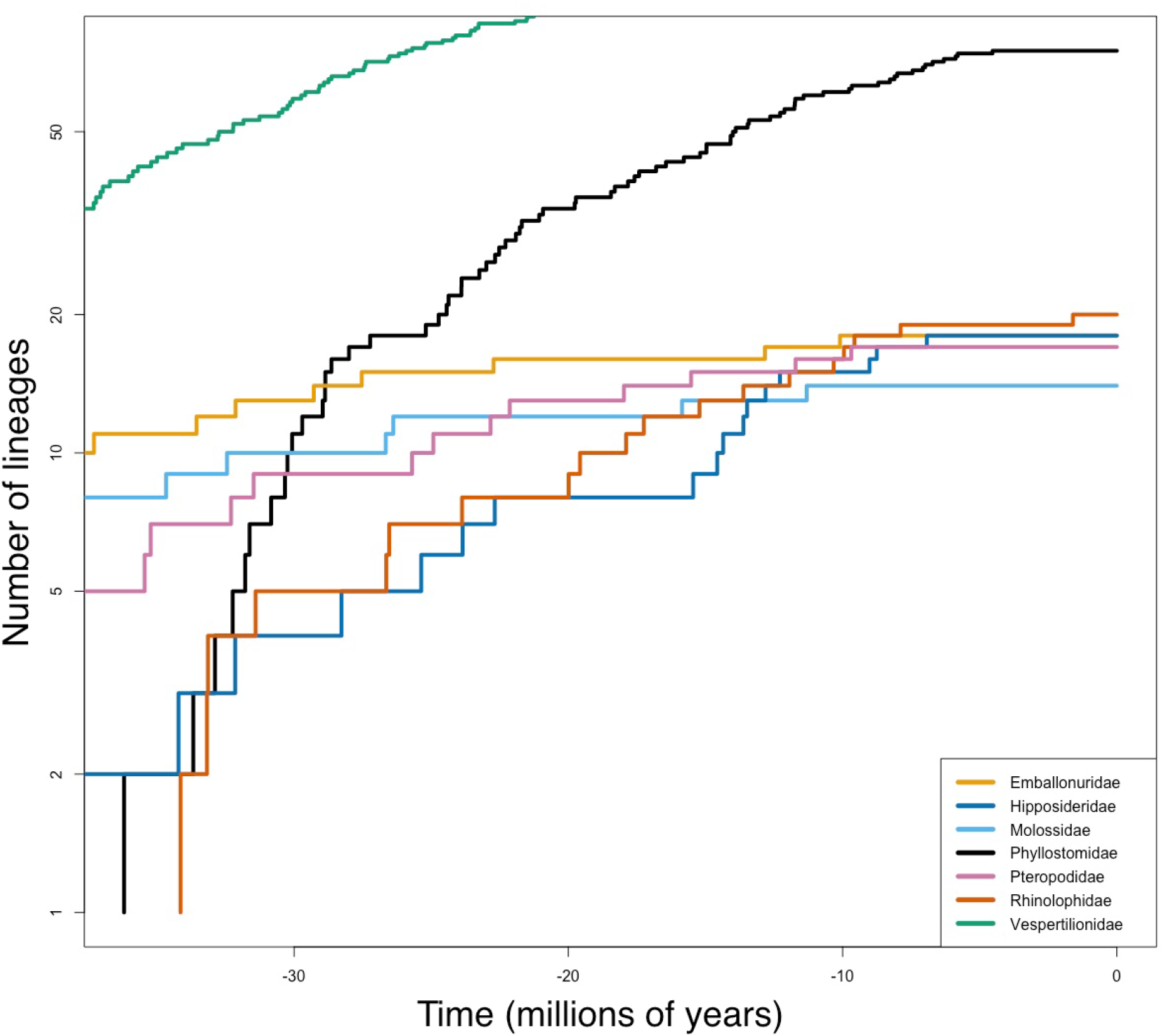
Lineage through time (LTT) plot based on the dated maximum likelihood RaxML-NG COI tree showing lineage accumulation in the last 30 million years for seven bat families. Plot scaled using the family Phyllostomidae (New World leaf-nosed bats, black line) as reference.

### Phylogenetic diversity and predictive modeling

Average MNTD and MPD values for the 25-eco dataset across the ecoregions were 0.38 and 0.62, respectively. There were no obvious global geographic patterns in PD as measured by MPD, except that the southern hemisphere contained higher levels of MPD (Fig. S4). Mideast Africa and Southeast Asia had the highest levels of MPD, followed by regions in Southern North America, Central America, Northern South America, and Southern Africa. Western North America had the lowest levels of MPD. Average PD values for the 109-eco dataset for MNTD and MPD across the ecoregions were 0.42 and 0.60, respectively. Ecoregions in Asia had the highest MPD values, followed by some areas in Africa (Fig. 1). Similarly, the southern, more tropical regions of the globe had higher levels of MPD.

MNTD as the response variable had low predictive power. For the 25-eco set, at most, 11.2% of the variation was explained by LGM variables. For the 109-eco set, at most, 19.9% of the variation was explained by geographic variables. Geography, the LGM, and climate change were the only groups of variables that produced models with predictive power for MNTD, though still relatively low. The models using MPD as a response variable explained anywhere from 47.4-67.8% [25-eco] or 33.5-58.4% [109-eco] of the variation depending on the model. Thus, we focus our attention on the results using MPD as our our response variable for predictive modeling (Table 1). While the percent variation explained differed among our MPD RF models, the geographic (25-eco: 67.8%; 109-eco: 57.4%) and climate change (25-eco: 65.3%; 109-eco: 42.2%) variables had the most explanatory power for both sets of ecoregions, though all categories had moderate predictive power. For each set of ecoregions, we conducted an RF analysis with the top 2-4 predictors from each category chosen based on the number of times the variable was in the top five predictors of the categorical RF analyses (Tables S8 & S9).

For the 25-eco set, there were 23 top variables in the final RF (Fig. 4) which included:

**Figure 4.**
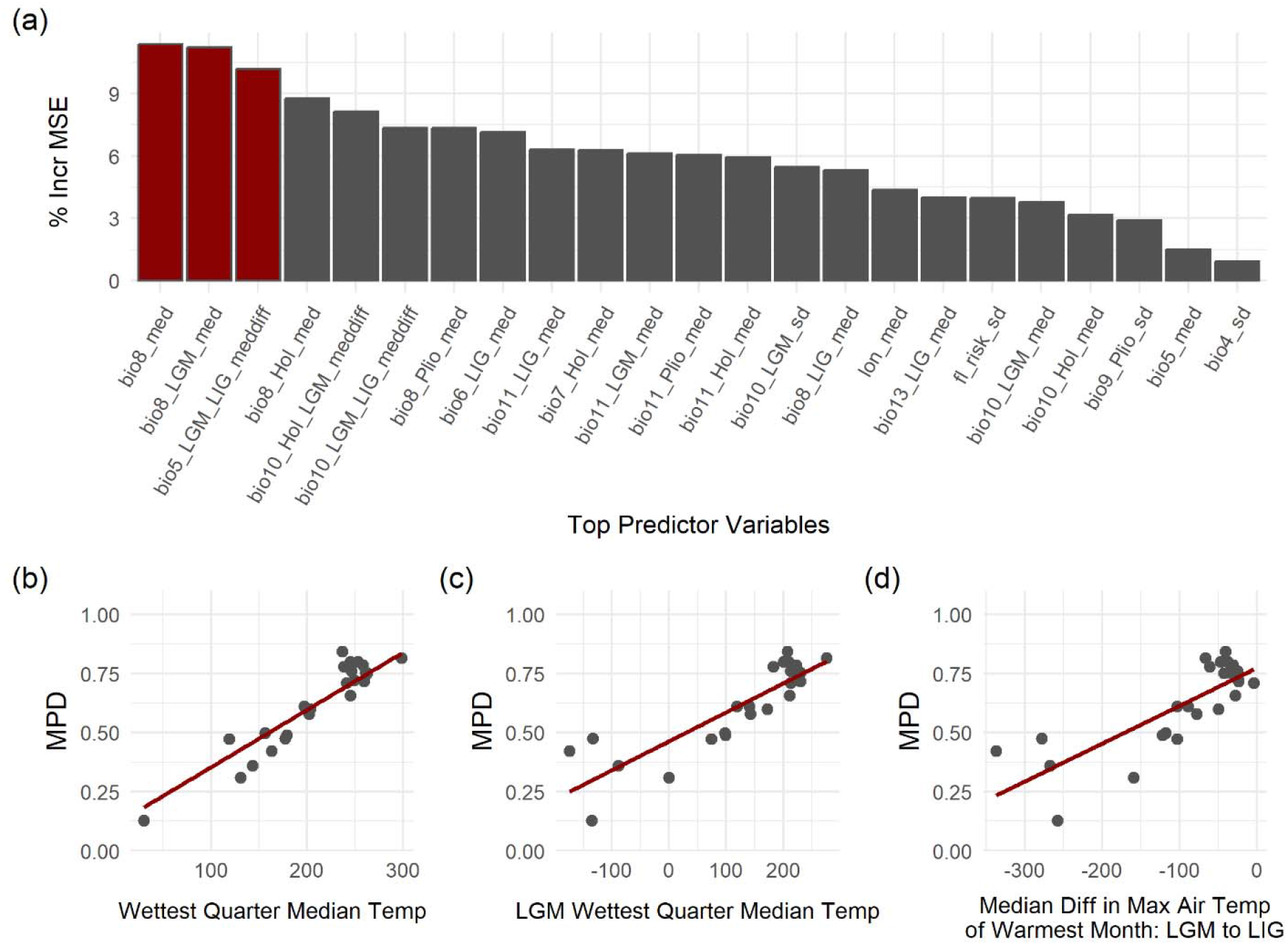
(a) Bar plot of % increase in MSE for all variables in the top random forest for the 25-ecoregion dataset. Red bars indicate those variables that were in the top five predictors 100% of the time for 100 random forest iterations. (b) Scatterplots with linear regression lines for the three top variables. All three were significant.

*Current BIOCLIM:* median temperature of the wettest quarter (bio8_med), median maximum temperature of the warmest month (bio5_med), and standard deviation of temperature seasonality (bio4_sd)

*Geography:* median longitude (lon_med) and standard deviation of flood risk(fl_risk_sd)

*Holocene BIOCLIM:* median temperature of the wettest quarter (bio8_Hol_med), median temperature of the coldest quarter (bio11_Hol_med), median temperature annual range (bio7_Hol_med), and median temperature of the warmest quarter (bio10_Hol_med)

*LGM BIOCLIM:* median temperature of the wettest quarter (bio8_LGM_med), median temperature of the coldest quarter (bio11_LGM_med), and standard deviation and median temperature of the warmest quarter (bio10_LGM_sd and bio10_LGM_sd)

*LIG BIOCLIM:* median minimum temperature of the coldest month (bio6_LIG_med), median temperature of the coldest quarter (bio11_LIG_med), median temperature of the wettest quarter (bio8_LIG_med), and median precipitation of the wettest month (bio13_LIG_med)

*Pliocene BIOCLIM:* median temperature of the wettest quarter (bio8_Plio_med), median temperature of the coldest quarter (bio11_Plio_med), and standard deviation of the mean temperature of the driest quarter (bio9_Plio_sd)

*Climate Change:* median maximum temperature of the warmest month between the LIG and LGM (bio5_LGM_LIG_meddiff), median temperature of the warmest quarter between the LIG and LGM (bio10_LGM_LIG_meddiff), and median temperature of the warmest quarter between the Holocene and LGM (bio10_Hol_LGM_meddiff).

This top variable RF model had high predictive power, explaining 78.91% of the variation in MPD. The top three variables from this analysis were current median temperature of the wettest quarter (bio8_med), LGM median temperature of the wettest quarter (bio8_LGM_med), and median maximum temperature of the warmest month between the LIG and LGM (bio5_LGM_LIG_meddiff), all showing up as top five predictors 100% of the time (Fig. 4). All three variables were significant after implementing a Bonferroni correction (0.05/3=0.0166) and accounting for spatial autocorrelation (Table 2).

**Table 2.** P-values for the top three variables in each of the three RF final models.

| Variable | eco-set | p-value |
| --- | --- | --- |
| median LGM BIO8 | 25 | <0.0001 |
| median present BIO8 | 25 | <0.0001 |
| median LGM-LIG BIO5 | 25 | <0.0001 |
| absolute latitude | 109 | 0.2654 |
| median gpw | 109 | 0.9779 |
| median BIO8 | 109 | <0.0001 |
| standard deviation kg2 | 41 | 0.0142 |
| standard deviation Pliocene BIO9 | 41 | 0.0138 |
| median LIG BIO11 | 41 | <0.0001 |

For the 109-eco set, there were 19 top variables in the final RF (Fig. 5) which included:

**Figure 5.**
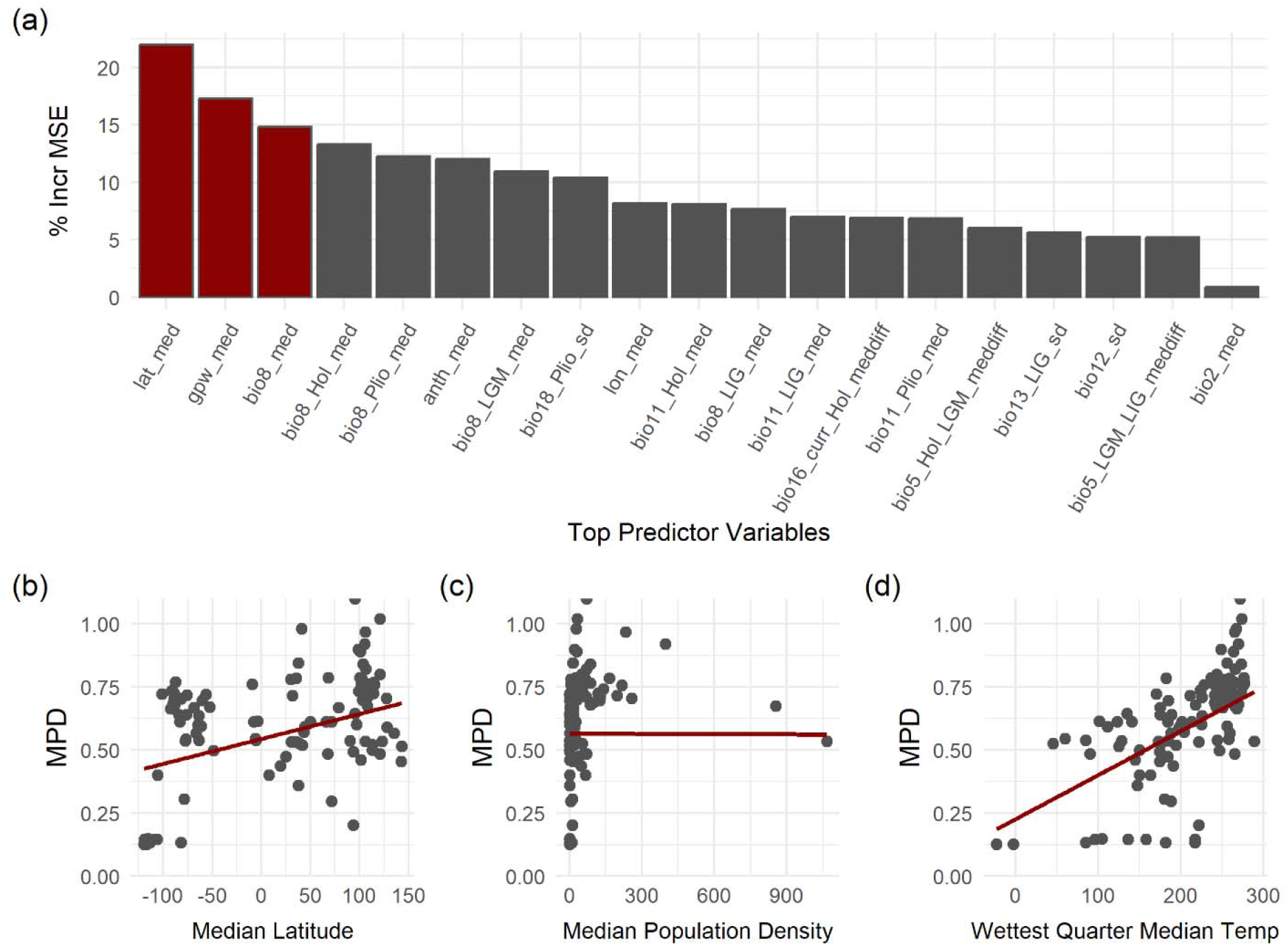
(a) Bar plot of % increase in MSE for all variables in the top random forest for the 109-ecoregion dataset. Red bars indicate those variables that were in the top five predictors 100% of the time for 100 random forest iterations. (b) Scatterplots with linear regression lines for the three top variables. Only median temperature of the wettest quarter was significant.

*Current BIOCLIM:* median temperature of the wettest quarter (bio8_med), median diurnal temperature range (bio2_med: mean of monthly [max temp – min temp]), and standard deviation of annual precipitation (bio12_sd)

*Geography:* median latitude (lat_med), median longitude (lon_med), median anthropogenic biomes (anth_med), and median SEDAC world population density (gpw_med)

*Holocene BIOCLIM:* median temperature of the wettest quarter (bio8_Hol_med) and median temperature of the coldest quarter (bio11_Hol_med)

*LGM BIOCLIM:* median temperature of the wettest quarter (bio8_LGM_med)

*LIG BIOCLIM:* standard deviation of the mean precipitation of the wettest month (bio13_LIG_sd), median temperature of the coldest quarter (bio11_LIG_med), and median temperature of the wettest quarter (bio8_LIG_med)

*Pliocene BIOCLIM:* standard deviation of mean precipitation of warmest quarter (bio18_Plio_sd), median temperature of the wettest quarter (bio8_Plio_med), and median temperature of the coldest quarter (bio11_Plio_med)

*Climate Change:* median maximum temperature of the warmest month between the Holocene and LGM (bio5_Hol_LGM_meddiff), median precipitation of wettest quarter between the present and Holocene (bio16_curr_Hol_meddif), and median maximum temperature of the warmest month (BIO5) between the LGM and LIG (bio5_LGM_LIG_meddiff).

This top variable RF model had high predictive power, explaining 56.78% of the variation in MPD. The top three variables from this analysis were median latitude (lat_med), median world population density (gpw_med), and current median temperature of the wettest quarter (bio8_med), all showing up as top five predictors 100% of the time (Fig. 5). Only current median temperature of the wettest quarter was statistically significant after implementing a Bonferroni correction (0.05/3=0.0166) and accounting for spatial autocorrelation (Table 2).

Finally, for the 109-eco set reduced to 41 ecoregions after removing PD estimates that were not significant after randomization, there were 18 top variables in the final RF (Fig. 6) which included:

**Figure 6.**
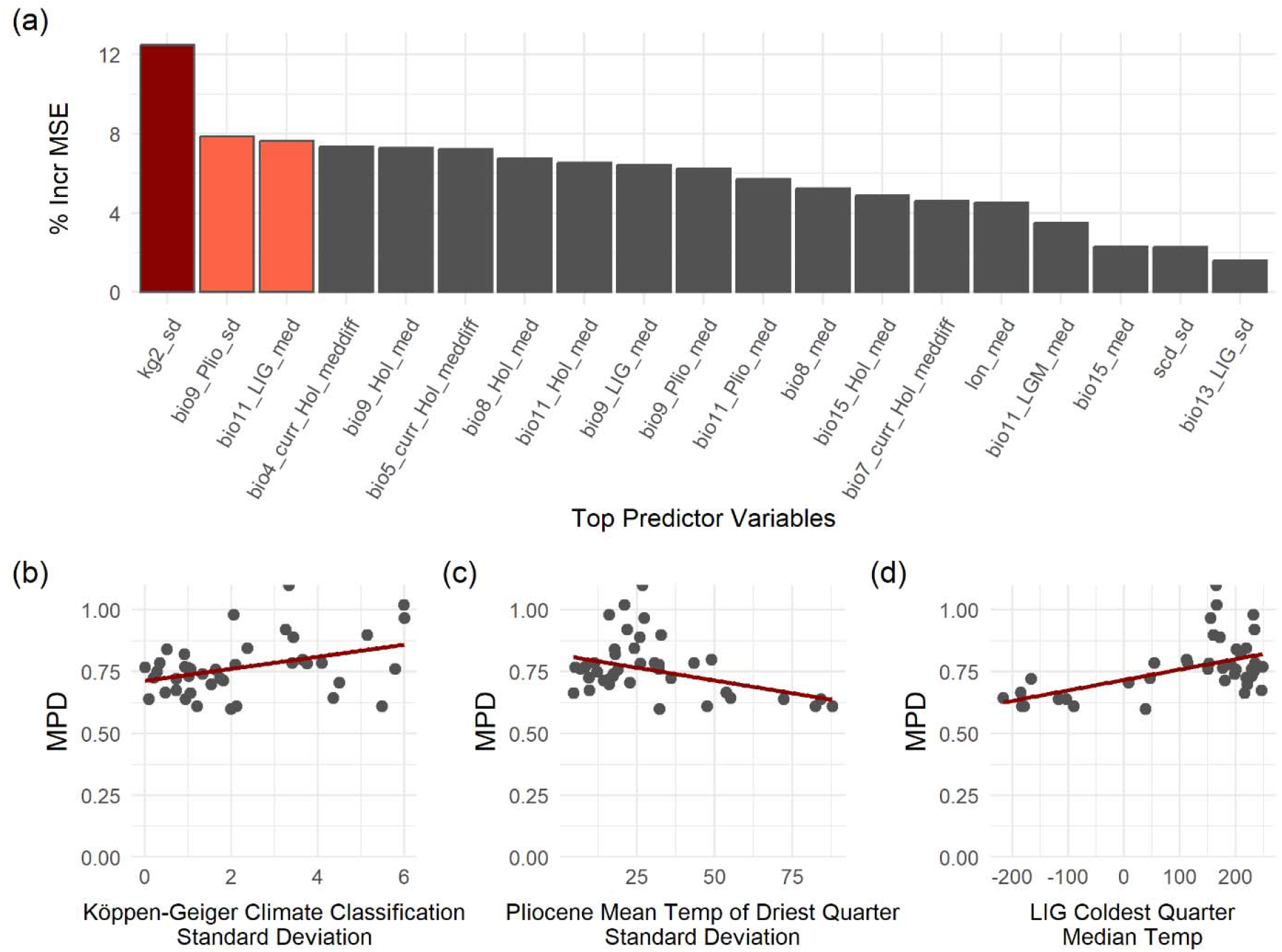
(a) Bar plot of % increase in MSE for all variables in the top random forest for the ecoregion dataset with only significant mpd values. Red bars indicate those variables that were in the top five predictors 100% of the time for 100 random forest iterations, and orange bars indicate those that were in the top five 75% or more of the time for 100 random forest iterations. (b) Scatterplots with linear regression lines for the three top variables. All three were significant.

*Current BIOCLIM:* median precipitation seasonality (bio15_med) and median temperature of the wettest quarter (bio8_med)

*Geography:* standard deviation of the Köpppen-Geiger climate classification (kg2_sd) and standard deviation of snow cover days (scd_sd)

*Holocene BIOCLIM:* median temperature of the driest quarter (bio9_Hol_med), median temperature of the coldest quarter (bio11_Hol_med), median precipitation seasonality (bio15_Hol_med), and median temperature of the wettest quarter (bio8_Hol_med)

*LGM BIOCLIM:* median temperature of the coldest quarter (bio11_LGM_med)

*LIG BIOCLIM:* median temperature of the coldest quarter (bio11_LIG_med), median temperature of the driest quarter (bio9_LIG_med), and standard deviation of the mean precipitation of the wettest month (bio13_LIG_sd)

*Pliocene BIOCLIM:* median temperature of the driest quarter (bio9_Plio_med), median temperature of the coldest quarter (bio11_Plio_med), and standard deviation of the mean temperature of the driest quarter (bio9_Plio_sd)

*Climate Change:* median maximum temperature of the warmest month between the present and Holocene (bio5_curr_Hol_meddiff), median temperature seasonality between the present and Holocene (bio4_curr_Hol_meddiff), and median temperature annual range between the present and Holocene (bio7_curr_Hol_meddiff)

This top variable RF model had similar predictive power to the full 109-eco set, explaining 47.03% of the variation in MPD. The top three variables from this model were standard deviation of the Köpppen-Geiger climate classification, Pliocene standard deviation of the mean temperature of the driest quarter (bio9_Plio_sd), and LIG median temperature of the coldest quarter (bio11_LIG_med). While only Köpppen-Geiger climate classification showed up as a top five predictor 100% of the time, the other two were in the top five 82% and 79% of the time. All were significant after implementing a Bonferroni correction (0.05/3=0.0166) and accounting for spatial autocorrelation (Table 2).

## Discussion

Terrestrial ecosystems have been rapidly altered by anthropogenic pressures (Ellis et al., 2021). The degradation and fragmentation of habitats and overexploitation of species globally have culminated in the current biodiversity crisis. Various conservation initiatives have proposed targets of 30 to 50% protection of the terrestrial biosphere to mitigate species loss (Dinerstein et al., 2019; Watson & Venter, 2017). Deciphering the environmental and spatial factors that shape different aspects of biodiversity is imperative to resolve regional and global patterns of diversity and can support conservation practices by identifying high levels of biodiversity that are less likely to be lost given future climate change. We leveraged publicly available, single-locus genetic data and spatial layers to estimate OTUs and examine the distribution of bat phylogenetic diversity (PD) at a global scale and its driving factors using random forest predictive modeling with a large set of starting predictor variables.

We produced a global bat phylogeny and defined OTUs using the Assemble Species by Automatic Partitioning (ASAP) method. While ASAP genetic groups generally matched current bat taxonomy, we encountered various groups in the dataset where merging and/or splitting of nominal species was common. These groups fit into different categories. For example, some merged groups included OTUs within genera with cryptic species that have undergone recent taxonomic revisions (e.g., Jarrín-V & Kunz, 2008; Parlos, et al., 2014; Velazco, et al., 2017; Calderón-Acevedo & Muchhala, 2018; Calderón-Acevedo & Muchhala, 2020; Calahorra-Oliart, et al., 2021; Calderón-Acevedo et al., 2022; Calahorra-Oliart et al., 2021; da Silva Fonseca et al. 2024; Mônico & Soto-Centeno 2024), particularly small Stenodermatine frugivores (Rodríguez-Posada, et al., 2018; Morales-Martínez, et al., 2021). Other mismatched groups included widely distributed insectivores that occur from Africa to Southeast Asia (Foley et al., 2015; Foley, et al., 2017; Patterson et al., 2019; Taylor et al., 2012; Csorba, et al., 2011). These examples highlight some of the many recent efforts to provide more stable taxonomy based on species delimitation (Calahorra-Oliart et al., 2021; da Silva Fonseca et al. 2024; Mônico & Soto-Centeno 2024). For example, the *Vampyressa melissa* species complex was recently split into multiple species with the use of *Cytochrome b* sequence data (Morales-Martínez et al., 2021). Three of these species, that are now considered at-risk due to their small ranges and rare occurrences, would have otherwise gone undetected without the use of genetic data, increasing their risk of extinction. Our ASAP delimitation results (Table S11) provide a guide for future similar taxonomic work in multiple clades that might contain cryptic species whose identification could have downstream conservation implications (Hending, 2024).

Patterns in OTU PD match those found in species richness gradients and global phylogenetic diversity patterns in birds (Udy et al., 2021; Barrios & Martinez-Nuñez, 2024). Southeast Asia is known for its high levels of bat species richness, and our analysis revealed that it also exhibits some of the highest levels of PD. Despite having the second highest bat species richness, the Neotropics showed lower phylogenetic diversity than parts of Africa and the Himalayas, both of which had the next highest levels of PD in our analysis. This matches previously found patterns in species diversity and other measures of PD in bats (López-Aguirre, et al., 2018). The fact that the Himalayas had relatively higher levels of PD but lower species richness, a pattern observed previously, might be particularly important because of this region’s high sensitivity to climate change (Chakravarty, et al., 2021). Furthermore, in birds, the only other group of flying vertebrates, PD was higher in mountainous regions while being lower in overall species richness (Jarzyna, et al., 2021; Montaño-Centellas et al., 2023). Similar to the latitudinal gradient of species diversity, we detected relatively high PD around the tropics compared to temperate areas (Fig. 1 & S4), and latitude was identified as an important predictor. Eastern South America, central and southern Africa, central Asia, northern Mexico, and southern Australia have been identified as places to expand protected areas as they are locations that have intact landscapes and transition potential (Dobrowski et al. 2021). Some of these regions overlap with the same areas where we detected high levels of PD in bats and could serve as important hotspots for future biodiversity conservation. The ecoregions we have identified with high PD using *COI* can be used as a starting point for local conservation efforts that want to incorporate PD and climate indicators in their decision making. However, it is important to note that overall sampling was lower in the global south, including large areas of South America and parts of central Africa, where more of our ecoregions did not have robust estimates of PD (according to SES resampling). With this caveat, the distribution of PD does not follow the same pattern as sampling (Fig. 1) and temperature variables and bioclimatic history consistently come out as top predictors even when ecoregions with weak PD estimates were removed from the analysis.

At both spatial scales, the 25- and 109-ecoregion sets, median temperature of the wettest quarter was a significant contributor to PD in bats and was moderately important when using the 41-eco set. Median temperature of the wettest quarter during the LGM and the change in median temperature of the warmest month between the LGM and LIG were also significant when looking at larger spatial units (25-eco set). Both the standard deviation of temperature of the driest quarter during the Pliocene and median temperature of coldest quarter during the LIG were significant when looking at smaller spatial units (41-eco set). These results are congruent with similar studies analyzing the relationship between temperature and various measures of phylogenetic diversity in bats and birds. Phylogenetic diversity measures like MNTD and MPD have been associated with temperature when looking at both Mexican bat communities and global bird diversity (Grimshaw & Higgins, 2017; Montaño-Centellas et al., 2023; Barrios & Martinez-Nuñez, 2024). Warm climates will result in more habitat stability and heterogeneity allowing for time and space to fill a diverse set of evolutionary niches (Kozak and Wiens 2010). Overall, temperature is a consistent factor that contributes to phylogenetic diversity. Additionally, the standard deviation of the Köppen-Geiger climate classification was important and significant. This further supports that habitat heterogeneity plays an important role in PD and should be considered when identifying areas for protection (Udy et al. 2021).

There are also indications that considering historical components is important for explaining PD (Montaño-Centellas et al., 2023; Paz et al., 2021; Svenning et al., 2015), and these variables are often not included in studies of biodiversity. Many of the temperature variables identified as top predictors in our analyses have a historical component (e.g. Mid-Holocene, LGM, LIG, and Pliocene). This could be key to understanding patterns of diversity given that climate oscillations during the Quaternary (i.e., Pleistocene and Holocene) have been associated with extinction, shaping genetic structure, and distribution of many small mammals in temperate regions (Blois, et al., 2010; Shafer, et al., 2010). Changes in climate during this time likely also played a significant role in the historical demography of bats (Carstens et al., 2018), conceivably contributing to niche evolution. We did detect the change in temperature from the LIG to LGM as important, potentially suggesting that while climate stability increases species diversity, change over time might influence PD differently. Braga et al. (2023) found that climate stability since the LGM increases relatedness in bat communities. It is possible that we did not identify climate change as a stronger predictor because its impact may vary and could be difficult to detect at a global scale (Kissling et al., 2021). Despite that, both deeper and recent climate seem to shape PD. The warmer Pliocene climate was a driver of speciation in African birds (Voelker, et al., 2010). However, we observed a negative relationship between temperature during the Pliocene and PD, suggesting that extremely hot and arid climates can limit speciation and niche evolution in bats. This information can be considered when making future predictions for the stability of species richness in bats over long timeframes, especially when the goal is to preserve PD.

We showed a slowdown in speciation rates over time in bat families globally. This pattern has been previously observed in studies sampling all bat families (Shi & Rabosky 2015) and in the superfamily Noctilionoidea (Rojas et al. 2016). These studies attributed speciation slowdowns to a combination of macroecological and biogeographic processes with protracted speciation after a diversification event related to trophic niche availability (Shi & Raboski 2015; Rojas et al. 2016). Generally, bats depict a saturated expansion scenario where species diversification plateaus towards the present, with Phyllostomidae showing a group-specific diversification pattern where species accumulation slowdown follows a more recent rapid rate of expansion. While we used a narrower genetic dataset compared to other studies, it still captured the signature of diversification scenarios observed across all bats. Importantly, the association between PD and climate that we estimated likely describes the observed patterns of bat evolutionary history. In our analysis, the apparent decoupling of PD estimates from the timing of increased speciation rate, for example in phyllostomids, highlights the importance of lineage specific factors that account for patterns at the ecoregion level and are independent of bursts in speciation rate. For example, Neotropical plants show a pattern of more frequent diversity expansions compared to tetrapods (Meseguer et al., 2022). These increases in diversification rate were stimulated by climatic cycles and the expansion of different biomes (e.g. Dick & Pennington, 2019). The increased plant diversification and resulting habitat heterogeneity associated with climate may explain the maintenance of higher regional PD in other organisms such as bats.

The measure of phylogenetic diversity being used also seems to influence which predictors are important. In contrast to analyses using MNTD and MPD, studies looking at Faith’s index saw precipitation variables as top predictors (Grimshaw & Higgins, 2017; Paz et al., 2021). We did not consider Faith’s index in our predictive modeling because it was highly correlated with OTU richness in our dataset, and we did not want our results to be driven by OTU richness. Although, we did detect some variation in predicting PD at different spatial scales. While the 25-eco worldwide dataset had top variables that were primarily temperature related, similar to analyses from smaller regions like the Atlantic Forest of Brazil and Mexico (Grimshaw & Higgins, 2017; Paz et al., 2021), the 109-eco data also identified median latitude and population density as important, though these were not significant when accounting for spatial autocorrelation. We believe it could be worthwhile to more systematically explore if the same patterns apply within and between ecoregions and what differences arise. For example, like our results, López⍰Baucells et al. (2022) found that at small spatial scales environmental variables do not seem to explain PD in aerial insectivorous bats, but PD was higher in larger forest fragments. This could influence how we think about conservation practices for species at different spatial scales in conjunction with evolutionary history, as it is not unusual for some patterns, like latitudinal diversity gradients or human impact, to be more or less strong at different spatial scales (Hillebrand, 2004; Barrios et al. 2024). Furthermore, protected areas do not always maintain within-species genetic diversity (Thompson, et al., 2021) or fully capture phylogenetic diversity (Saraiva, et al., 2018). Future studies at local scales should examine the usefulness of protected areas for preserving phylogenetic diversity in bats (e.g., Leon-Alvarado & Miranda-Esquivel, 2023). Specifically, protected areas could be evaluated on how much PD they currently have and what the future projections of their climate would look like for variables that have been deemed important (i.e., temperature).

Monitoring biodiversity is essential for conserving current and future levels of biodiversity. In addition to the thousands of phylogenetic and phylogeographic studies that produce mtDNA sequence data, environmental DNA metabarcoding and barcoding efforts produce a large amount of sequence data that can be used for these purposes (Rodríguez-Ezpeleta et al. 2021). When it comes to biodiversity assessments needed to justify and inform conservation efforts, we have shown that open-source, single-locus data can be a valuable tool. While our study analyzed environmental and geographic predictors of bats, this computationally efficient method could be applied to any taxa of interest with sufficient georeferenced data and used to inform conservation efforts that prioritize the evolutionary history of species. Both phylogenetic and functional diversity have been found to predict biodiversity effects like biomass production (Flynn, et al., 2011). Since collecting functional diversity data is more resource intensive, focusing on PD may be a more efficient and effective way of documenting biodiversity and ecosystem services that go beyond species richness (Molina-Venegas, et al., 2021; Forest, et al., 2007; Pio, et al., 2011). PD has also been shown to be connected to the stability of vegetation production, which is an important factor in assessing an environment’s ability to withstand climatic changes (Mazzochini, et al., 2019).

Global bat phylogenetic diversity is highest around the tropics of Asia, Central America, South America, and Africa which aligns well with patterns of species richness, but assessments at different spatial scales may provide slightly different results. However, using a broad-scale ecoregion approach, studies of phylogenetic diversity in other taxa can provide areas of overlap among organisms that could make for the most valuable targets for conservation. In bats, phylogenetic diversity is best explained by temperature variables while also considering a historical component. By understanding what environmental and geographic factors foster higher levels of phylogenetic diversity, we can better determine what environments contain the diversity necessary to withstand climate change and where conservation resources will have the largest impact. PD also has more evolutionary potential and can be useful to consider for conservation purposes long-term. While this information would never be a sole determinant of conservation focus areas, the analysis is easy to run and can be an additional decision-making tool when it comes to allocation of conservation resources.

## Acknowledgements

Funding provided by National Science Foundation DBI-1911293 to TAP. We thank Radford University’s Office of Undergraduate Research for supporting AEG through the Highlander Research Rookies program.

## Data availability statement

All intermediate datafiles and list of DNA sequence accessions are available as supplemental tables (Table S1-8). All data analysis scripts and supplemental files 1-12 are on Dryad: Reviewer URL: http://datadryad.org/stash/share/Wrp3Bem5LbdGpxC8bdzp8h0f3eyMJW0zoc6hNW11y8A.

