## Additional Supplemental Information for "Environmental and geographic drivers of global bat phylogenetic diversity"

**Supporting information for: Environmental and geographic drivers of global bat phylogenetic diversity**

**List of Tables**

**Table S1**. Accession numbers for all *COI* sequences. Those that were removed after data cleaning steps are documented. *PhylogatR* IDs include GenBank accessions followed by GBIF accession, or a set of zeros followed by BOLD accessions.

**Table S2**. Breakdown of merged Resolve ecoregions with DNA sequences present that were included in Random Forest analyses to create the eco-25 dataset.

**Table S3**. Data layers used as predictor variables with links. Those with missing data and not included in Random Forest are indicated.

**Table S4**. Final dataset used as input for Random Forest analyses 25-eco. Includes PD values, number of OTUs per ecoregion, and values for all predictor variables.

**Table S5**. Final dataset used as input for Random Forest analyses 109-eco. Includes PD values, number of OTUs per ecoregion, and values for all predictor variables.

**Table S6.** Predictor variable pairwise correlations for eco-25 dataset.

**Table S7.** Predictor variable pairwise correlations for eco-109 dataset.

**Table S8**. Variable importance and counts for the number of times they are top variables in 100 iterations of random forest for the eco-25 dataset.

**Table S9**. Variable importance and counts for the number of times they are top variables in 100 iterations of random forest for the eco-109 dataset.

**Table S10**. Variable importance and counts for the number of times they are top variables in 100 iterations of random forest for the eco-41 dataset.

**Table S11**. ASAP groupings for all DNA sequences. Species are labeled as matched (all individual sequences from the species only belonged to one ASAP group), merged (individual sequences from the species were in the same ASAP group as another species), or split (individual sequences from the species were found in multiple ASAP groups).

**List of Supporting Files**

Supporting file S1 Nucleotide alignment COI_OTUs_withSP_final.fasta

Supporting file S2 Best Tree COI_OTUs_withSP_final.fasta.raxml.bestTree

Supporting file S3 Family/Subfamily constraint file COI_OTUs_withSP_final_constraint

Supporting file S4 ML Trees COI_OTUs_withSP_final.fasta.raxml.mlTrees

Supporting file S5 Starting Tree COI_OTUs_withSP_final.fasta.raxml.startTree

Supporting file S6 Bootstrap support on Best Tree COI_OTUs_withSP_final.fasta.raxml.support

Supporting file S7 Bootstrap Trees COI_OTUs_withSP_final.fasta.raxml.bootstraps

Supporting file S8 Best Model COI_OTUs_withSP_final.fasta.raxml.bestModel

Supporting file S9 Rooted Best Tree rooted_COI_OTUs_withSP_final_bestTree.tree

Supporting file S10 TreePL Configuration File COI_OTUS_treePL

Supporting file S11 TreePL Dated Tree dated_besttree.tre

Supporting file S12 Superfamily constraint file COI_OTUs_withSP_final_Superfamily_constraint

**List of data analysis scripts**

**ASAP_seqs.R**

Randomly pulls one DNA sequence per ASAP group.

**bat_env_geo.R**

Calculates the mean and standard deviation of all geographic and climatic variables. Also calculates the difference in means between historical and present day BIOCLIM variables.

**bat_lm.R**

Runs linear regressions for top variables. Also includes code for linear regression scatterplots.

**env_corr.R**

Calculates correlation coefficients between all geographic and climatic variables.

**MAP.R**

Creates sequence count heatmaps.

**PD.R**

Calculates all 3 phylogenetic diversity measures (Faith’s Index, MNTD, MPD) for each ecoregion.

**phylogeny_reconstruction.txt**

Contains all IQ-Tree2, RAxML-NG, and TreePL commands used for analysis.

**quantify_ASAP.R**

Determines which species are matched, merged, and split based on ASAP results.

**RegionPHeatMap.R**

Creates a raster file with heatmap of MPD values.

**summry_sp_delim.R**

Summarizes species delimitation results. (Table S8 without matched, merged, split columns)

**variable_reduction_RF.R**

Reduces variable list based on correlation cutoffs, categorizes variables, and runs all random forest analyses.

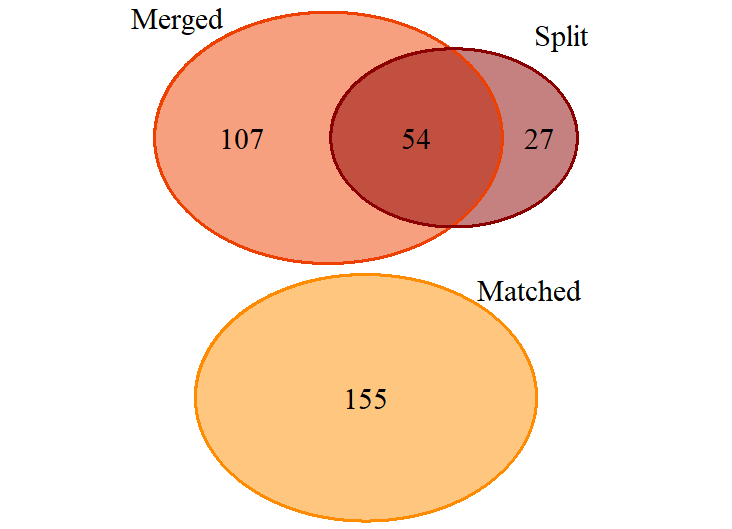

**Figure S1**. ASAP results. Visual breakdown of the matched, merged, and split designations given to all species in our data based on the ASAP analysis. See Table S8 description for details on matched, merged, and split.

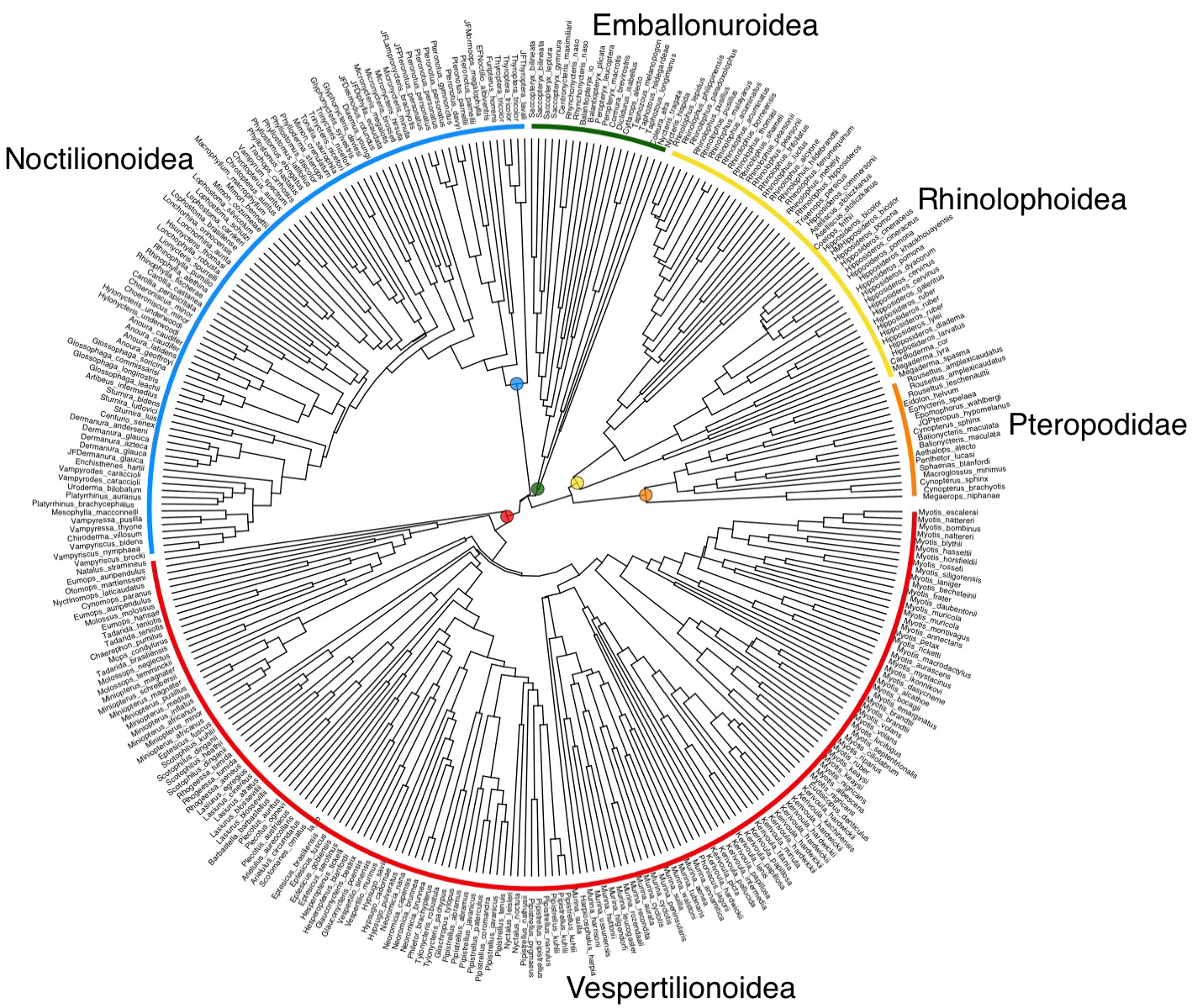

**Figure S2**. Maximum likelihood phylogeny including 17 bat families worldwide produced using COI sequences. Tips represent species labels. Family and superfamily labels are included as reference following the time-scaled phylogeny in Fig. 2.

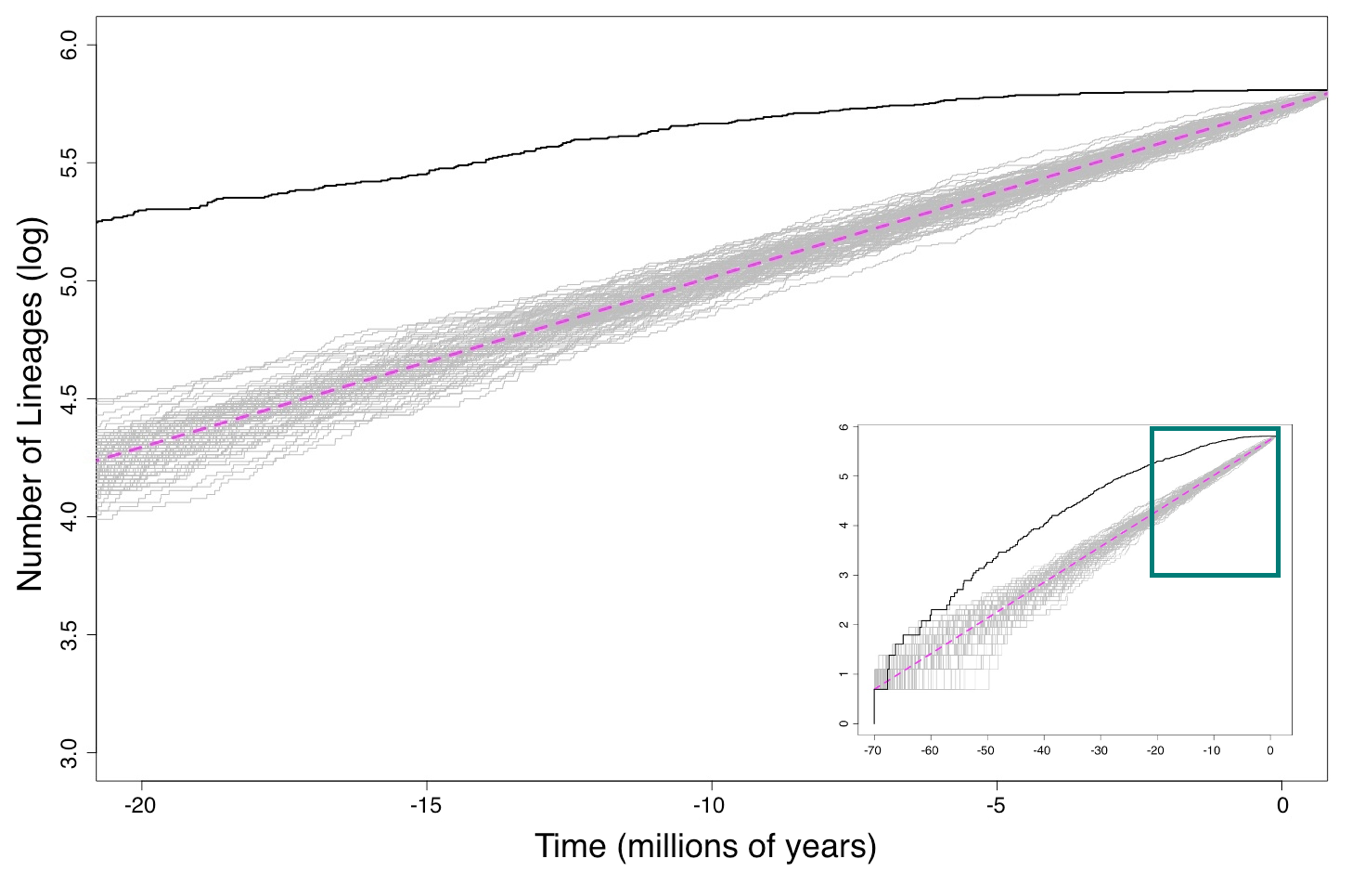

**Figure S3**. Lineage through time (LTT) plot based on the dated maximum likelihood RaxML-ng COI tree showing lineage accumulation in the last 20 million years for all 17 bat families combined. Inset shows the full LTT plot. Black line represents the empirical LTT estimated from the phylogeny. Gray lines represent 100 simulated LTT plots using a birth-death process; trend of simulated species accumulation shown in magenta, to depict expected species accumulation curves for an expected phylogeny of comparable sampling size. Y-axis converted to log scale to aid in visualization.

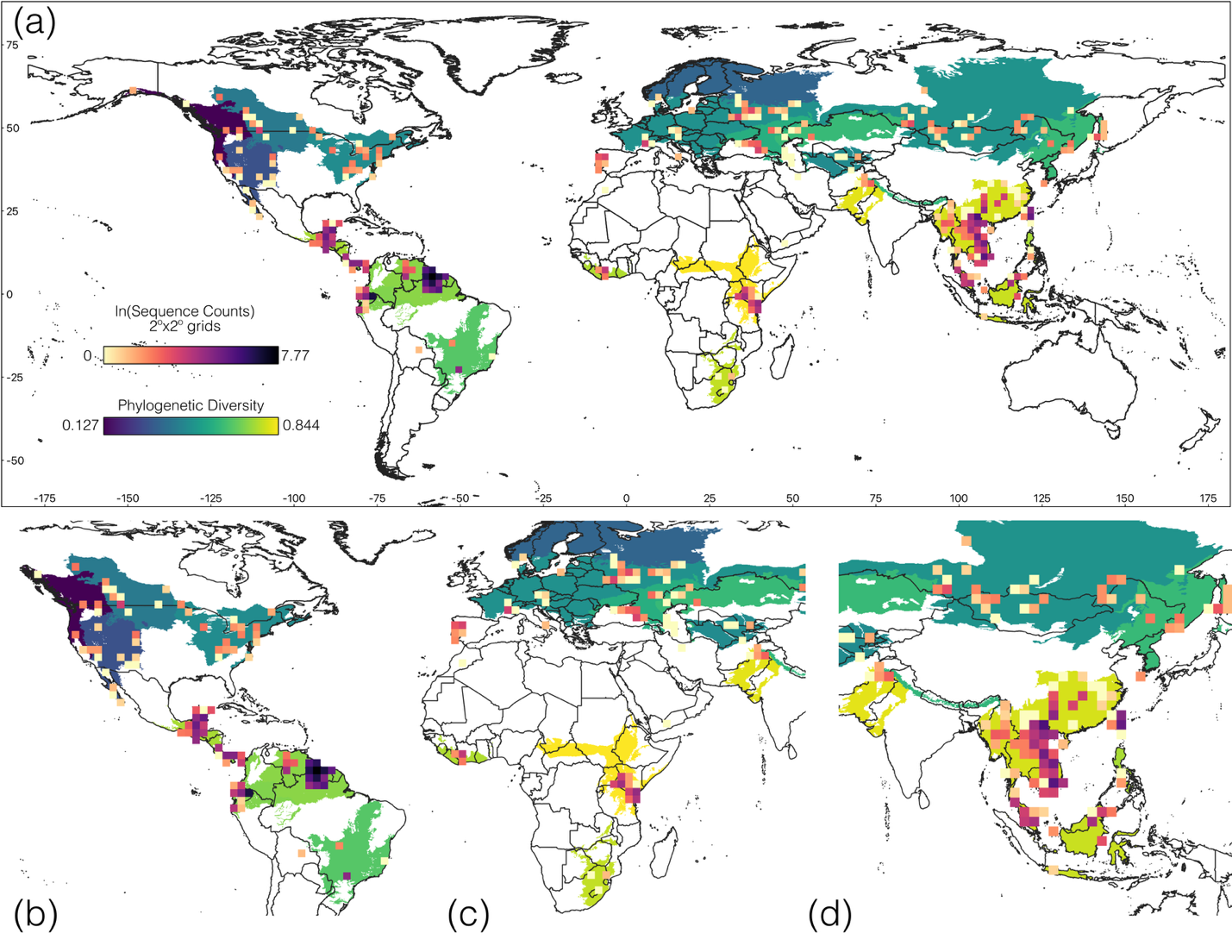

**Figure S4**: 25-ecoregions. (a) Phylogenetic diversity of bats Worldwide and number of sequences (given in natural logarithm of actual sequence numbers in a 2x2 degree grid, for better visualization). (b) The Neotropics (c) East Africa (d) Southeast Asia
